# Multi-Structure Segmentation of Type B Aortic Dissection from CT Images with U-Net

**DOI:** 10.64898/2026.11.08.765191

**Authors:** Ruonan Gong, Jiasong Chen, Linchen Qian, Yuhang Du, Bradley G. Leshnower, Minliang Liu, Liang Liang

## Abstract

Type B aortic dissection (TBAD) is a life-threatening condition in which a tear in the intimal layer separates the aortic wall into a true and a false lumen. Assessment of the lumen, the intimal flap, and false-lumen thrombosis from CT images is important for risk analysis and treatment planning. This work applied U-Net-based deep neural networks to the automatic segmentation of these structures in TBAD. We collected 46 CT scans from 20 patients, with voxel-wise annotations of three structures: the aortic lumen, the intimal flap, and the thrombus. We explored four models: two convolutional networks, nnU-Net (2D and 3D), and two transformer-based architectures, TransUNet (2D and 3D).

We observed that (1) for both nnU-Net and TransUNet, the 3D models outperformed their 2D counterparts on all three structures. The difference was small for the aortic lumen but large for the thin flap and the thrombus. The best model, nnU-Net 3D, reached a DSC of 93.05% for the lumen, 50.22% for the flap, and 69.94% for the thrombus. (2) The intimal flap was the most difficult structure for all models. Even the best 3D models showed large thrombus boundary errors in a few scans. These results provide a baseline for developing more advanced models for accurate segmentation of these structures in TBAD.

## 1 Introduction

Cardiovascular disease remains a leading cause of death worldwide [1]. Aortic dissection is an acute aortic syndrome in which blood enters the media through an intimal tear and creates a false lumen separated from the true lumen by the intimal flap. According to the Stanford classification, dissections that do not involve the ascending aorta are classified as type B [2]. Morphological features such as false-lumen diameter and volume, the extent of the dissection flap, and the degree of false-lumen thrombosis have been associated with disease progression and adverse outcomes [3].

Computed tomography angiography (CTA) is the standard imaging modality for diagnosis and surveillance of TBAD. Extracting these morphological features requires segmenting the aortic lumen, the intimal flap, and the thrombus in 3D, which is time-consuming per scan when done manually. Therefore, automatic segmentation is of considerable clinical interest.

Machine learning, particularly deep learning, has recently become a powerful tool in medical image analysis. Deep neural networks can learn hierarchical image representations directly from annotated data, and they have reached expert-level performance in tasks such as the segmentation of organs at risk in head and neck CT for radiotherapy planning [4]. For image segmentation, the U-Net [5] combines a contracting path that captures context with an expanding path that recovers spatial detail through skip connections, and its volumetric extension [6] processes 3D image patches directly, which allows the network to exploit inter-slice context in CT and MRI. However, convolutions only process a local neighborhood at a time, which makes it difficult to capture long-range dependencies in the image. Vision transformers [7] address this with self-attention, which relates all parts of the image to each other. In medical image segmentation, they have been used either as the encoder [8] or combined with CNNs in hybrid designs [9].

Deep learning has also been applied to TBAD. Cao et al. [10] used 3D convolutional networks to segment the whole aorta, the true lumen, and the false lumen from CT scans, and Chen et al. [11] improved the segmentation by first simplifying the aortic anatomy in a multi-stage framework. More recently, nnU-Net-based pipelines have been used to identify and segment aortic dissection automatically [12]. Public datasets have made such work easier to compare: ImageTBAD [13] provides annotations of the true lumen, the false lumen, and false-lumen thrombus, and Mayer et al. [14] released expert annotations of the true and false lumen. However, most of these works focus on the true and false lumen. Thrombus is not annotated in some datasets [14] and is merged with the lumen in some methods [15], and the intimal flap is rarely segmented as a structure of its own.

nnU-Net [16] is a self-configuring framework that automatically adapts preprocessing, network topology, and training to a given dataset. It has been a strong baseline across many segmentation challenges, including the recent AortaSeg24 challenge on multi-class aortic segmentation, where most participating teams built on it [17].

In this work, we aim to develop deep learning models that can accurately segment the aortic lumen, the intimal flap, and the thrombus in TBAD from CT scans. We explore U-Net-based architectures for this task, including convolutional (nnU-Net) and transformer-based (TransUNet) networks operating on 2D slices or 3D volumes. We then evaluate their segmentation quality with overlap and boundary metrics.

## 2 Materials and Methods

### 2.1 Datasets

Our dataset consists of 46 CT scans from 20 patients with TBAD. Example scans and annotations are shown in Figure 1. All data were de-identified. Several patients were imaged at multiple time points (1–4 scans per patient). Each scan was treated as a separate case, and all scans of a patient were always kept in the same data split (more details in Section 3.2). The median in-plane voxel spacing is 0.80 mm (range 0.33–0.98 mm) and the median slice spacing is 1.25 mm (range 0.33–2.50 mm). The median image size is 512 *×* 512 *×* 446 voxels. Three structures were annotated:

**Figure 1:**
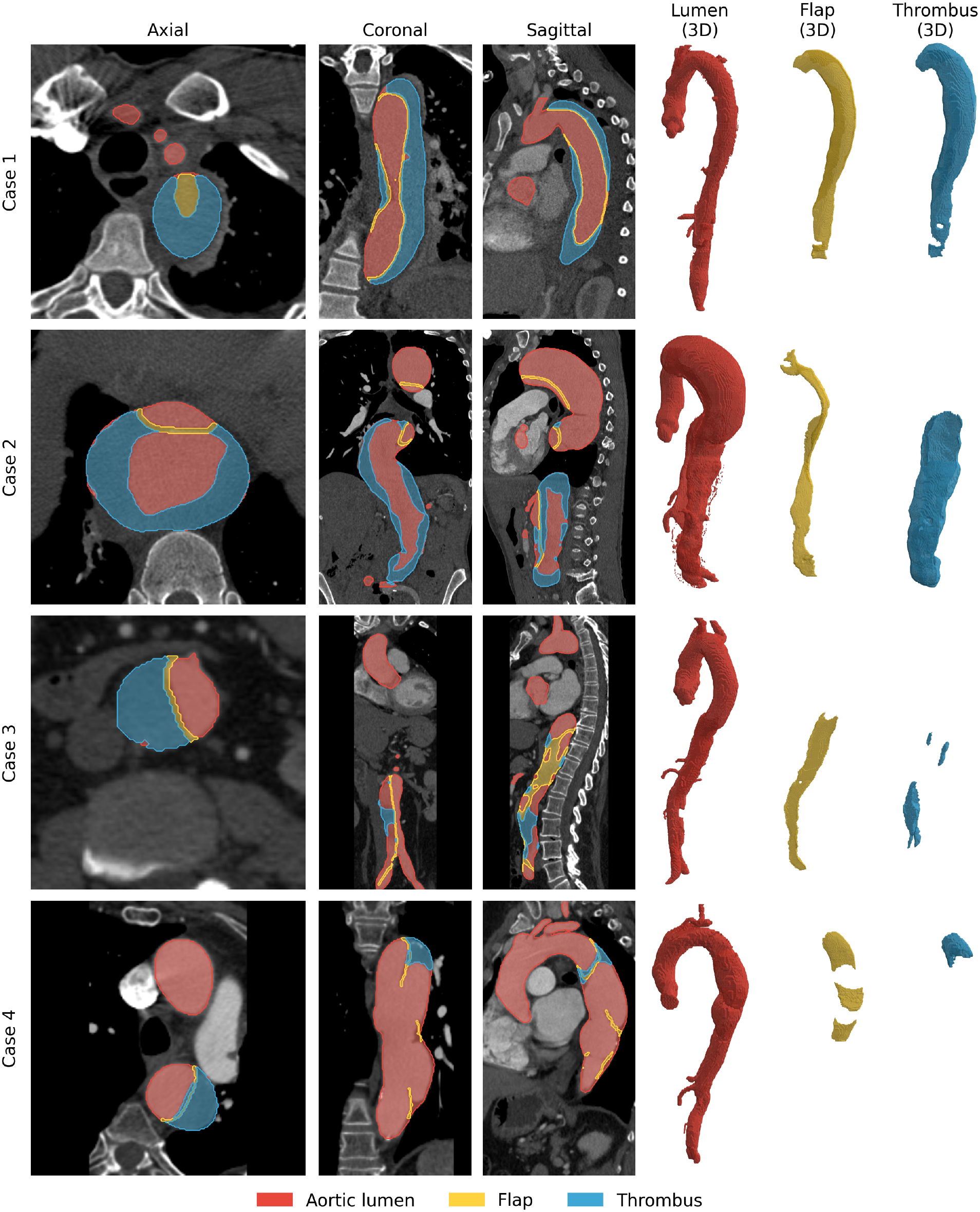
Example CT images and annotations from four patients. From left to right: axial, coronal, and sagittal slices with the annotations overlaid, and 3D renderings of the aortic lumen (red), the intimal flap (yellow), and the thrombus (blue).

- **Aortic lumen** (label 1): the contrast-filled lumen of the aorta, i.e. the true lumen and the perfused part of the false lumen.
- **Intimal flap** (label 2): the dissection membrane separating the true and false lumen.
- **Thrombus** (label 3): thrombosed parts of the false lumen.

### 2.2 Network Structures

**nnU-Net** nnU-Net [16] is a self-configuring segmentation framework based on the U-Net architecture. It uses fixed rules to configure preprocessing, network topology, patch size, and batch size. We used nnU-Net v2 with the residual encoder preset (ResEnc L) [18] and trained both its 3D full-resolution and its 2D configuration. The 3D network, shown in Figure 2a, has six resolution stages with 3 *×* 3 *×* 3 convolutions, and the number of feature maps grows from 32 to 320. It operates on patches of 192 *×* 192 *×* 192 voxels at a target spacing of 0.80 *×* 0.80 *×* 1.25 mm. The 2D network in Figure 2b has eight resolution stages with 3 *×* 3 convolutions and 32 to 512 feature maps, and segments each axial slice independently on patches of 512 *×* 512 pixels at an in-plane spacing of 0.80 *×* 0.80 mm.

**Figure 2:**
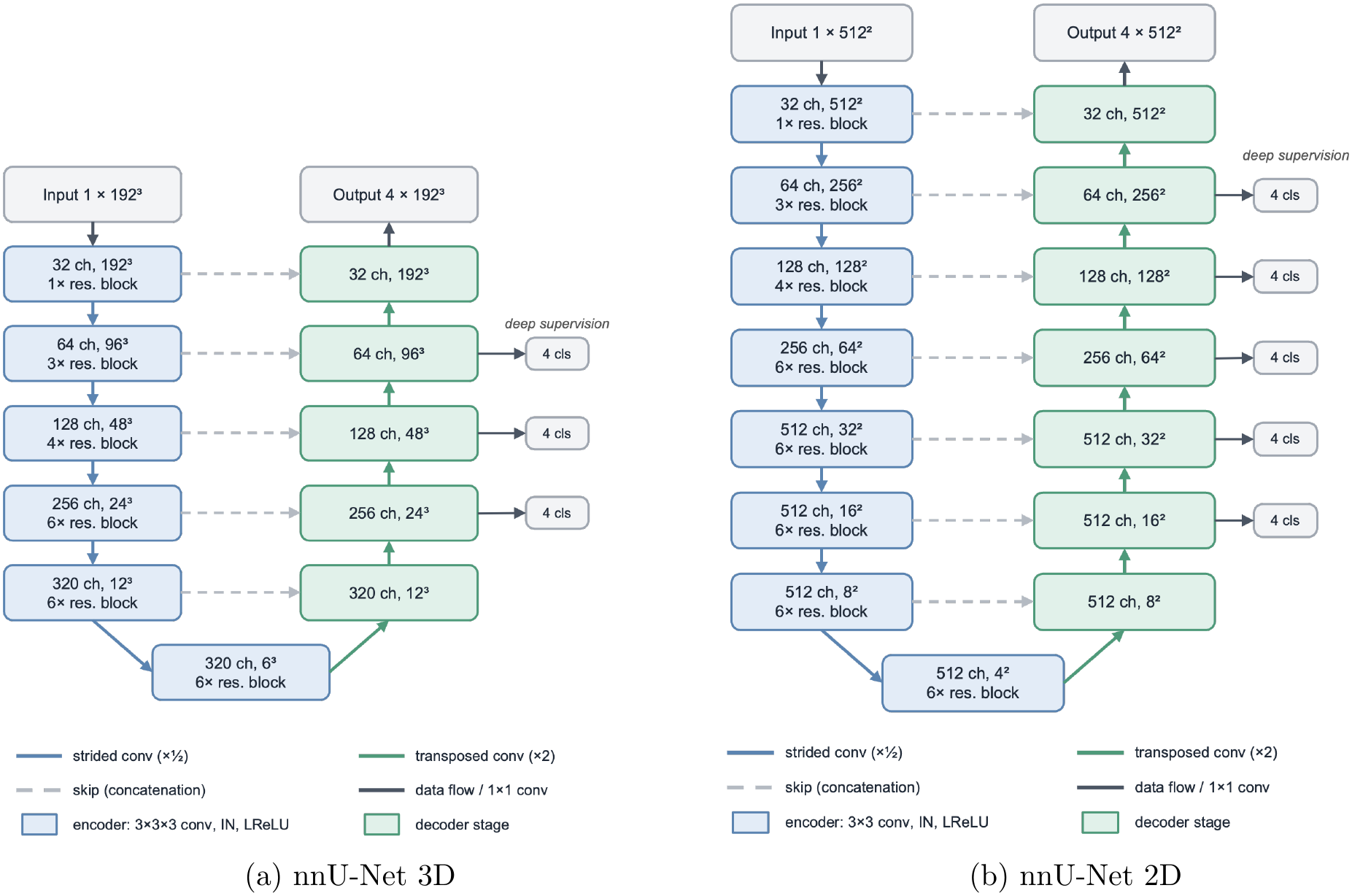
nnU-Net architectures as configured for our dataset. Each box gives the number of feature channels and the spatial size of a resolution stage.

#### TransUNet

TransUNet [9] is a hybrid CNN-transformer encoder-decoder model. The 2D version, shown in Figure 3b, processes each axial slice independently. Its encoder is a ResNet-50 followed by a 12-layer Vision Transformer (ViT-B/16) [7] and is initialized with weights pretrained on ImageNet-21k. The decoder upsamples the transformer output and combines it with the ResNet features through skip connections. TransUNet 3D [19] extends TransUNet to volumetric data within the nnU-Net (v1) framework, as shown in Figure 3a. The encoder is a 3D CNN followed by a 12-layer transformer, which is initialized with the same pretrained ViT-B/16 weights. The decoder is based on Mask2Former [20]. It uses a set of learnable queries, and each query predicts the mask of one structure.

**Figure 3:**
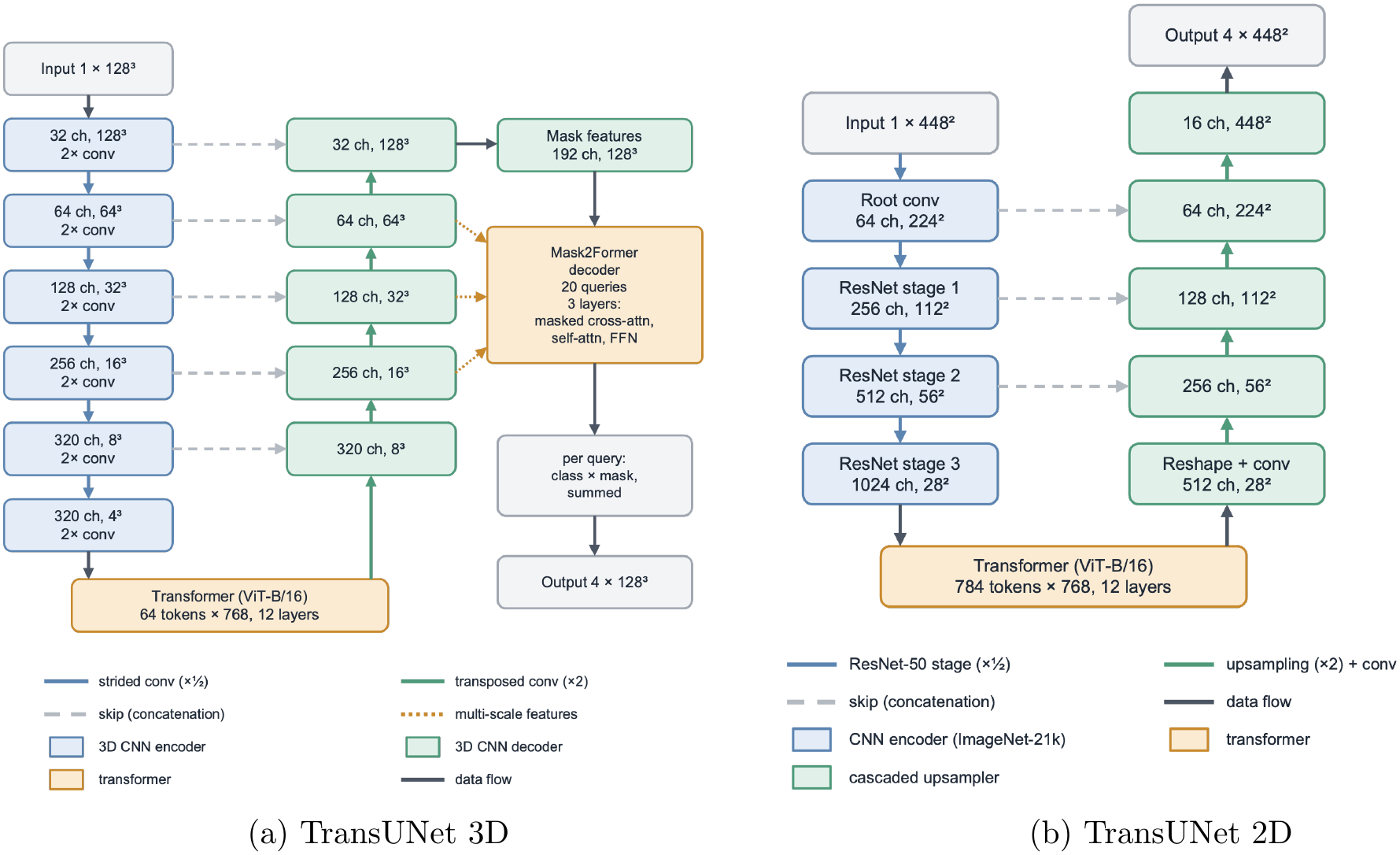
TransUNet architectures. (a) The 3D encoder+decoder variant combines a 3D CNN encoder, a transformer bottleneck, and a Mask2Former-based query decoder. (b) The 2D model combines a ResNet-50 encoder, a 12-layer Vision Transformer, and a cascaded upsampler.

### 2.3 Loss Functions

nnU-Net (2D and 3D) and TransUNet 2D were trained with a combination of Dice loss and cross-entropy loss,

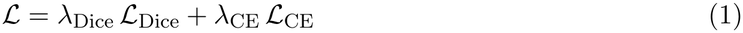

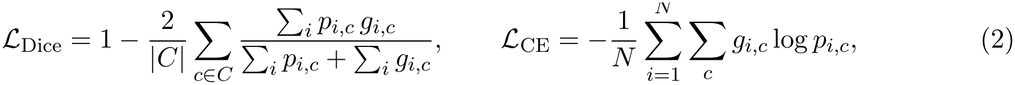

where *p*_*i,c*_ is the predicted softmax probability of class *c* at voxel *i, g*_*i,c*_ is the one-hot ground truth, and *N* the number of voxels. nnU-Net uses *λ*_CE_ = *λ*_Dice_ = 1 and computes the Dice term over the foreground classes only. It applies deep supervision, which computes the loss at several decoder resolutions with weights halving at each lower resolution. TransUNet 2D uses *λ*_CE_ = *λ*_Dice_ = 0.5 and includes the background class in the Dice term.

TransUNet 3D follows the set-prediction formulation of Mask2Former [20, 19]. Each class present in a training patch defines one target mask, and the 20 queries are assigned to these targets by Hungarian matching with the cost 2 *C*_cls_ + 5 *C*_BCE_ + 5 *C*_Dice_. For each decoder output, the loss is

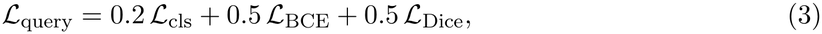

where ℒ_cls_ is the cross-entropy of the query class predictions, and *L*_BCE_ and *L*_Dice_ are the binary cross-entropy and Dice losses between the masks of the matched queries and their targets. The final loss averages the loss of the last decoder layer with the mean loss of the intermediate decoder layers, and an additional Dice loss on lower-resolution outputs of the CNN decoder provides deep supervision.

## 3 Experiments and Results

### 3.1 Data Processing

For nnU-Net (2D and 3D) and TransUNet 3D, the images were preprocessed with the standard nnU-Net pipeline v2 and v1 respectively. Each scan was cropped to the non-zero region, resampled to a common target spacing, and its CT intensities were clipped to the 0.5th–99.5th percentiles of the foreground and normalized to zero mean and unit variance.

For TransUNet 2D, CT intensities were clipped to [−100, 600] HU and scaled to [0, 1]. Slices were resampled in-plane to 0.8 mm, and scans with a slice spacing below 0.8 mm were resampled to 0.8 mm along the slice direction so that no single scan dominates the training set. Each slice was then cropped or padded to 448 *×* 448 pixels around the body center, so that the aorta has a similar physical size in all scans.

For all models, the predictions were mapped back to the original CT grid before evaluation.

### 3.2 Experiment Setup

All models were evaluated with five-fold cross-validation on the 46 scans. The folds were defined at the patient level, so that all scans of a patient were always in the same fold, and the same five splits were used for all models. In each fold, the scans of 16 patients were used for training and the scans of the remaining four patients for validation, which gave 36 or 37 training scans and 9 or 10 validation scans per fold. Each scan was therefore segmented exactly once by a model that had not seen it during training, and all metrics were computed on these out-of-fold predictions.

nnU-Net was trained with its default settings: 1000 epochs of 250 iterations, stochastic gradient descent (SGD) with Nesterov momentum, an initial learning rate of 0.01 with polynomial decay, and extensive data augmentation. The 3D network used patches of 192 *×* 192 *×* 192 voxels with a batch size of 2, and the 2D network patches of 512 *×* 512 pixels with a batch size of 35.

TransUNet 2D was trained with the default settings: 150 epochs with SGD, an initial learning rate of 0.01, and a batch size of 24, starting from ImageNet-21k pretrained weights. TransUNet 3D was trained for 1000 epochs of 250 iterations on 128 *×* 128 *×* 128 patches with a batch size of 2 and an initial learning rate of 0.01 with warm-up and cosine decay. Its transformer was initialized with the same pretrained ViT weights. No post-processing was applied to any model.

All experiments were run on NVIDIA H200 GPUs, with one GPU per fold.

### 3.3 Evaluation Metrics

All metrics were computed per scan and per structure at the original CT resolution. With TP, FP, FN, and TN denoting the true positive, false positive, false negative, and true negative voxels of a structure, the overlap metrics are

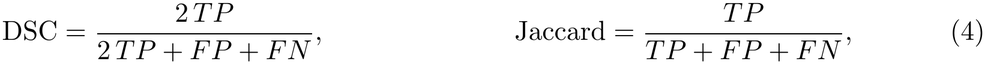

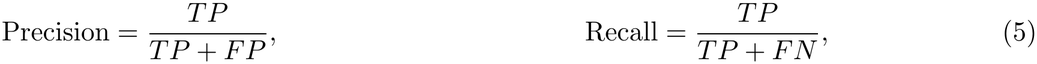

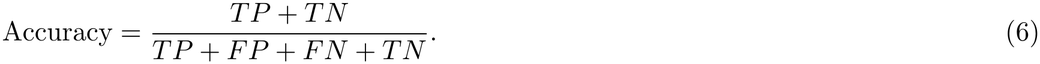

Boundary metrics are computed from the surface voxels *S*_*P*_ and *S*_*G*_ of the prediction and the ground truth, using Euclidean distances in millimeters:

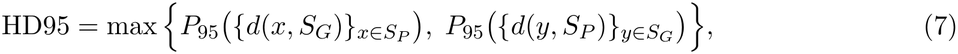

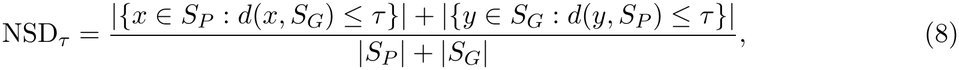

where *d*(*x, S*) = min_*y*∈*S*_∥*x* − *y*∥, *P*_95_ denotes the 95th percentile, and *τ* is the distance tolerance, which we set to 1 mm.

In addition, we computed the Chamfer distance (CD) between surface meshes. Each binary structure mask was converted to a triangular surface mesh with the marching cubes algorithm, using the voxel spacing so that vertex coordinates are in millimeters. The CD is

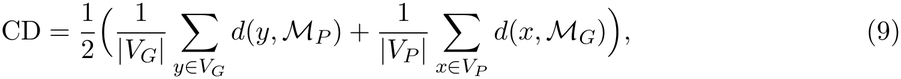

where ℳ_*P*_ and ℳ_*G*_ denote the prediction and the ground truth meshes. *V*_*P*_ and *V*_*G*_ are their vertices, and *d*(*x*, ℳ) is the distance from a vertex *x* to the closest point on the surface mesh ℳ.

### 3.4 Experiment Results

Table 1 reports the results for the individual structures, and Table 2 for the whole aorta. Figure 4 shows the predictions of all four models for two scans.

**Table 1:** Per-structure results, 5-fold cross-validation (mean *±* SD over scans, and HD95 and CD additionally with the median in brackets). The best mean value of each metric is shown in bold. CD: Chamfer distance between surface meshes.

| Structure | Metric | nnU-Net 3D | TransUNet 3D | nnU-Net 2D | TransUNet 2D |
|---|---|---|---|---|---|
| Aortic lumen | DSC (%) | **93.05**±1.96 | 92.59±2.54 | 89.81±6.49 | 88.87±4.48 |
|  | HD95 (mm) | **5.71**±4.66 [4.40] | 10.84±17.61 [4.78] | 13.71±15.67 [9.13] | 16.30±20.11 [9.99] |
|  | CD (mm) | **1.11**±0.59 [0.94] | 1.44±1.15 [1.02] | 1.81±1.24 [1.45] | 2.29±1.89 [1.67] |
|  | NSD@1mm (%) | **78.03**±7.81 | 77.05±7.78 | 69.65±11.03 | 66.24±11.11 |
| Intimal flap | DSC (%) | **50.22**±14.77 | 49.76±14.29 | 43.15±15.18 | 40.88±16.26 |
|  | HD95 (mm) | 21.34±29.04 [7.02] | **16.69**±18.10 [9.02] | 27.78±28.42 [17.93] | 25.60±24.65 [13.87] |
|  | CD (mm) | 3.22±3.85 [1.54] | **2.70**±2.59 [1.92] | 4.58±4.99 [2.25] | 4.55±4.62 [2.37] |
|  | NSD@1mm (%) | **70.34**±17.51 | 69.03±17.02 | 62.03±18.70 | 58.63±20.30 |
| Thrombus | DSC (%) | **69.94**±22.86 | 66.29±26.06 | 52.71±25.19 | 50.79±23.46 |
|  | HD95 (mm) | **45.42**±70.71 [14.84] | 46.48±73.79 [14.93] | 65.79±73.07 [31.65] | 57.76±73.66 [26.15] |
|  | CD (mm) | 5.61±8.28 [2.40] | **5.37**±6.93 [2.57] | 10.54±13.47 [5.28] | 11.72±17.07 [5.46] |
|  | NSD@1mm (%) | **53.18**±17.45 | 52.49±15.65 | 36.58±16.36 | 34.46±15.11 |

**Table 2:** Segmentation performance for the whole aorta (lumen ∪ flap ∪ thrombus), 5-fold cross-validation, *n* = 46 scans. The best mean value of each metric is shown in bold.

|  | nnU-Net 3D |  |  | TransUNet 3D |  |  | nnU-Net 2D |  |  | TransUNet 2D |  |  |
|---|---|---|---|---|---|---|---|---|---|---|---|---|
|  | Min | Max | Mean±SD | Min | Max | Mean±SD | Min | Max | Mean±SD | Min | Max | Mean±SD |
| DSC (%) | 87.28 | 97.18 | **94.49**±1.95 | 86.49 | 97.11 | 93.91±2.57 | 53.94 | 96.23 | 89.90±6.68 | 69.93 | 95.31 | 89.00±5.67 |
| HD95 (mm) | 1.83 | 26.40 | **5.65**±5.15 | 1.82 | 100.02 | 10.73±18.66 | 2.84 | 65.87 | 14.74±13.82 | 3.86 | 123.36 | 17.88±20.19 |
| CD (mm) | 0.50 | 2.87 | **1.09**±0.57 | 0.59 | 7.30 | 1.47±1.29 | 0.82 | 8.11 | 2.20±1.52 | 0.88 | 12.19 | 2.69±2.19 |
| NSD@1mm (%) | 47.68 | 90.50 | **76.41**±10.49 | 46.53 | 90.05 | 74.92±10.31 | 26.70 | 83.95 | 62.76±14.34 | 26.00 | 78.62 | 59.84±13.83 |
| Jaccard (%) | 77.43 | 94.52 | **89.62**±3.41 | 76.19 | 94.39 | 88.62±4.42 | 36.93 | 92.74 | 82.20±9.29 | 53.76 | 91.04 | 80.61±8.66 |
| Precision (%) | 78.72 | 98.71 | **95.84**±3.20 | 78.16 | 98.44 | 95.33±3.87 | 81.40 | 99.53 | 95.23±4.02 | 76.84 | 99.06 | 94.34±4.84 |
| Recall (%) | 85.95 | 98.56 | **93.32**±3.07 | 81.09 | 98.22 | 92.71±3.63 | 36.99 | 97.14 | 86.08±10.45 | 54.08 | 97.13 | 85.06±9.77 |
| Accuracy (%) | 99.40 | 99.98 | **99.82**±0.20 | 99.33 | 99.98 | 99.81±0.21 | 98.09 | 99.97 | 99.67±0.41 | 98.93 | 99.97 | 99.66±0.36 |

**Figure 4:**
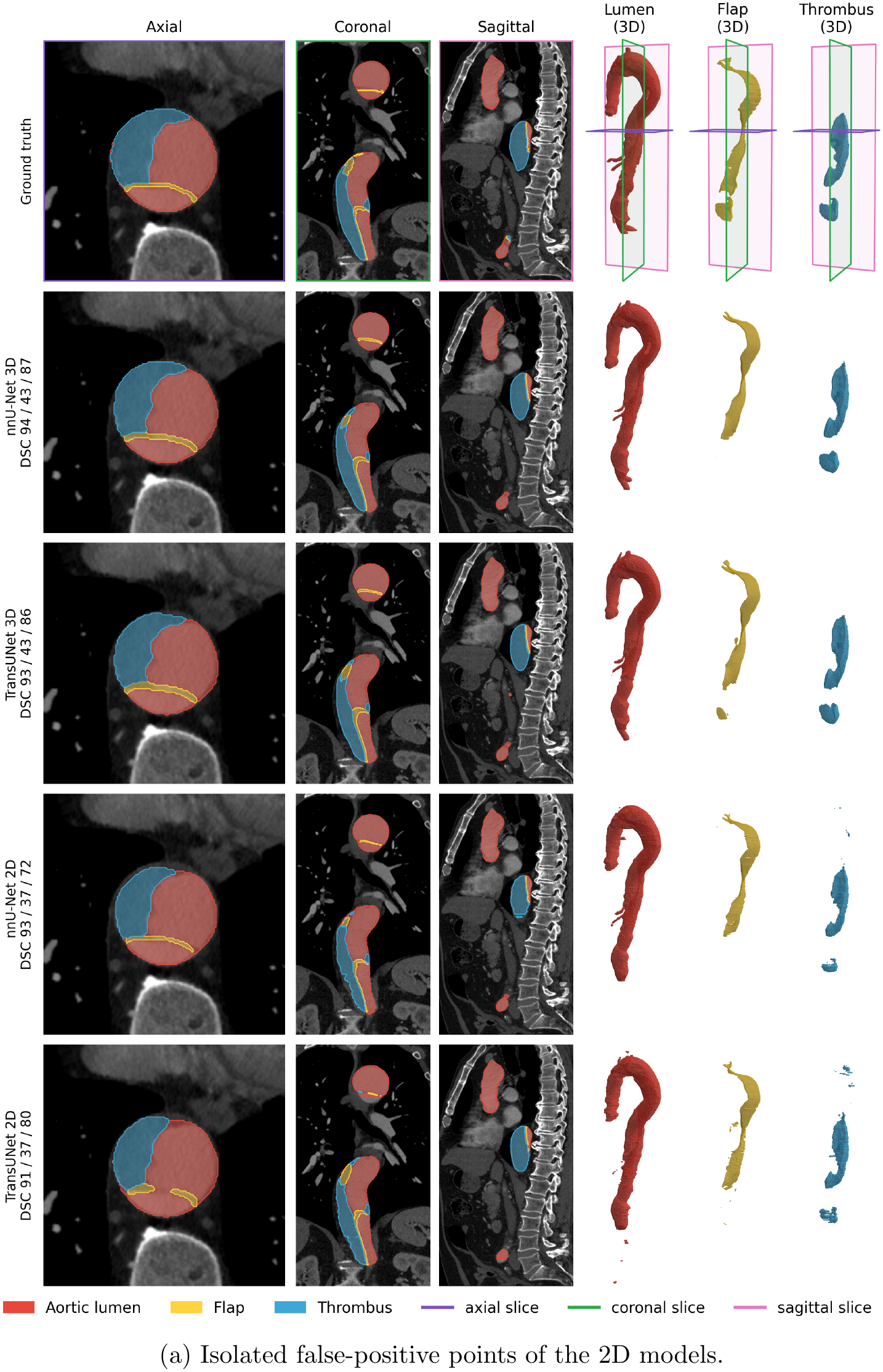
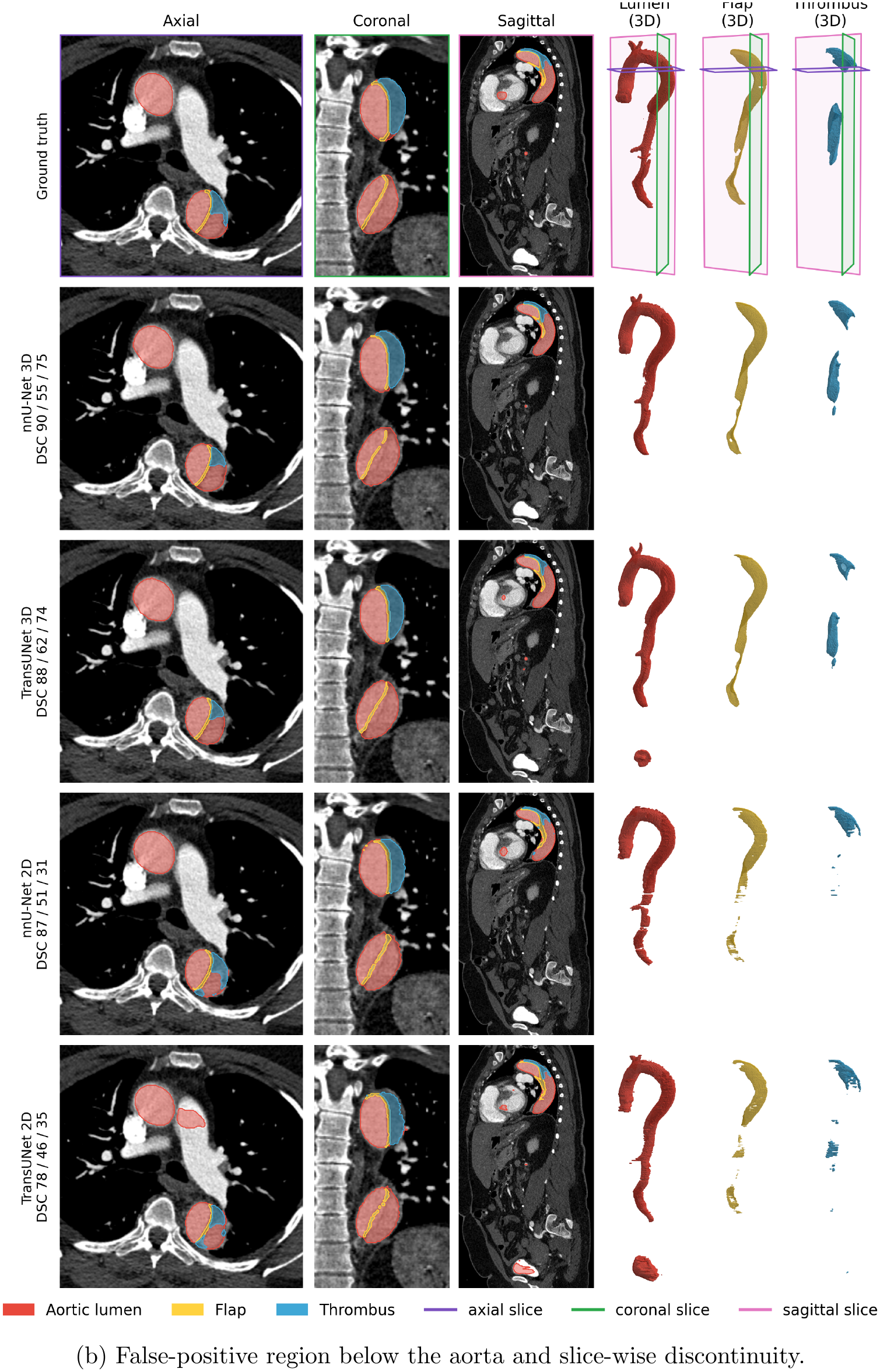
Predictions of all four models for two scans. In each panel, rows (top to bottom) show the ground truth and the predictions of nnU-Net 3D, TransUNet 3D, nnU-Net 2D, and TransUNet 2D, with the per-case DSC (%) of the lumen, flap, and thrombus. Columns show axial, coronal, and sagittal slices and 3D renderings of the aortic lumen, the flap, and the thrombus. The colored planes in the ground-truth renderings mark the slice positions. (a) nnU-Net 2D and TransUNet 2D produce scattered false-positive points outside the aorta, whereas the 3D models do not. (b) TransUNet 3D and TransUNet 2D predict a false-positive region below the aorta, and the prediction of nnU-Net 2D is discontinuous along the vessel, being present in some slices and missing in others. Predictions of all four models for a second scan.

As shown in Table 1, segmentation performance varied across the three structures. The aortic lumen achieved the highest DSC across all four models. The thrombus showed intermediate DSC values, and the intimal flap was the most challenging structure. In terms of model performance, nnU-Net achieved slightly higher DSC values than TransUNet for both 2D and 3D configurations across all three structures. For the whole aorta (Table 2), nnU-Net 3D and TransUNet 3D reached higher mean DSC than nnU-Net 2D and TransUNet 2D.

HD95 measures the boundary distance error between the predicted segmentation and the ground truth, with lower values indicating better boundary agreement. Among the three structures, the aortic lumen had the lowest HD95 values across the models, while the thrombus had the highest. This difference is also reflected in the Chamfer distance, which measures the average deviation between the predicted and reference surfaces. For the aortic lumen and whole aorta, the 3D models achieved an average surface deviation of approximately 1 mm. For the intimal flap and thrombus, the Chamfer distances were considerably lower than the corresponding HD95 values. This may suggest that although the average surface deviation was relatively small, larger errors were present in some local regions. Such errors may include missed portions of the thin flap or false-positive regions distant from the reference annotation. NSD provides a complementary measure of surface agreement by quantifying the proportion of surface points within a specified distance threshold. The aortic lumen had the highest NSD values across the four models, while the intimal flap also had relatively high NSD values despite its lower DSC. This indicates that a substantial proportion of the flap surface points remained within the 1 mm tolerance, even though the overall volumetric overlap was limited.

We observed two typical types of prediction errors. The first was discontinuity in the predicted aorta along the vessel. Compared with the 3D models, the 2D models showed similar precision but lower recall, indicating that they missed more portions of the aorta without substantially increasing predictions outside the reference region. A likely reason is that the 2D models segment each axial slice independently, so their predictions are not consistent between neighboring slices. In the coronal and sagittal views and in the 3D renderings shown in Figure 4b, this appears as a discontinuous prediction along the vessel, which is present in some slices and missing in others. In contrast, the 3D models, which process several slices at once, produced smooth and largely continuous surfaces.

A second type of error was isolated false-positive regions outside the aorta. As shown in Figure 4a, the 2D models produced scattered false-positive areas. Larger false-positive regions below the aorta occurred for both 2D and 3D models. Figure 4b shows such regions for TransUNet 2D and TransUNet 3D. Although such regions are small and hardly affect the DSC, they determine the HD95 of the affected scans. This may explain why the mean HD95 was much larger than the median for several models and structures, especially for thrombus.

Overall, with the same input dimensionality, nnU-Net and TransUNet performed similarly. nnU-Net 3D achieved the best mean value for most metrics, whereas TransUNet 3D achieved the lowest mean CD for the flap and thrombus and the lowest mean HD95 for the flap. For the flap and thrombus, nnU-Net 3D had the lower median Chamfer distance but a larger mean, i.e., it was more accurate on most scans but showed larger errors on a few of them. TransUNet 3D reached a similar median HD95 for the lumen as nnU-Net 3D but a larger mean, because of the distant false-positive regions described above.

## 4 Discussion and Conclusion

In this work, we investigated U-Net-based deep learning models for the segmentation of the aortic lumen, the flap, and the thrombus in TBAD from CT images. Four models were trained and evaluated with identical cross-validation splits: nnU-Net and TransUNet, each in a 2D and a 3D version. Generally, both nnU-Net and TransUNet achieved comparable performance when using the same input dimensionality, while the 3D configurations achieved better results than their 2D counterparts. The best results were obtained with nnU-Net 3D, closely followed by TransUNet 3D.

The advantage of the 3D models can be explained by the shape of the target. The aorta, the flap, and the thrombus are continuous structures that extend over hundreds of slices, and their appearance in one slice is strongly related to the neighboring slices. The 2D models segment each slice independently and cannot use this context. As a result, their predictions were not continuous along the vessel: parts of the aorta were missed in some slices, which lowered the recall, and scattered false-positive regions appeared outside the aorta, which increased the boundary errors. The 3D models produced smooth and continuous surfaces. The performance improvement from 3D context was small for the aortic lumen, which has a high contrast and is easy to segment even slice by slice, but large for the flap and the thrombus, whose identification depends more on the surrounding anatomy.

The flap was the most difficult structure for all models. The relatively high NSD values indicate that a substantial proportion of the predicted and reference surface points were within the 1 mm tolerance, despite the low DSC. This difference may be related to the thin and geometrically complex nature of the intimal flap, for which small boundary deviations can substantially affect volumetric overlap. The thrombus presented a different challenge. Its low contrast with the surrounding tissue and substantial variation in extent across patients may contribute to the large variability in its segmentation metrics.

A second source of error, shared by the 2D and the 3D models, was isolated false-positive regions far from the aorta. These regions are small and have little effect on the DSC, but they dominate the HD95 of the affected scans and explain why the mean HD95 was much larger than the median. Because such regions are spatially separated from the aorta, simple post-processing, for example keeping only the largest connected component, could remove many of them. We did not apply any post-processing in this study in order to compare the networks themselves.

In conclusion, 3D U-Net-based models segmented the aortic lumen, the intimal flap, and the thrombus in TBAD more accurately than their 2D counterparts, and nnU-Net 3D achieved the best overall performance. The thin flap and the low-contrast thrombus remain the main challenges, and the presented models provide a baseline for developing more accurate methods for these structures.

## Acknowledgments

This work was supported in part by the National Science Foundation under Grants DMS 2436630 and 2436629, and National Institutes of Health grant R01HL155537. We thank Kinza Johnson, Henry Blount, Marina Dominguez, and Caleb Dalton at Texas Tech University for their work on image annotation.

